# Visual physiology of Australian stingless bees

**DOI:** 10.1101/2024.11.14.623562

**Authors:** Bhavana Penmetcha, Laura A Ryan, Yuri Ogawa, Nathan S Hart, Ajay Narendra

## Abstract

Stingless bees engage in a range of visually guided behaviours that require relatively high spatial resolution and contrast sensitivity. Although the eyes of honeybees, bumblebees, carpenter bees, and sweat bees have been studied extensively, there is limited knowledge of stingless bees. Here, we studied two sympatric Australian species, *Tetragonula carbonaria* and *Austroplebeia australis*, which are important crop pollinators. The bigger *A. australis* had more and larger ommatidial facets compared to *T. carbonaria*. Using pattern electrophysiology, we showed that *A. australis* had higher contrast sensitivity (13.07) compared to *T. carbonaria* (5.99), but their spatial resolving power did not differ (0.53 cycles deg^-1^). We discuss these differences in visual physiology in the context of the distinct foraging behaviours of the two species.

## Introduction

Vision is a fundamental sensory modality used by insects for foraging, sexual selection, navigation and communication. Two significant visual capabilities for an insect are spatial resolving power and contrast sensitivity. Spatial resolving power is the ability to distinguish between small objects and resolve fine details in a scene. Contrast sensitivity is a measure of the ability to discriminate objects as their achromatic contrast decreases (O’Carroll and Wiederman 2014; Ogawa et al. 2019). These properties are dictated by the physiological and anatomical characteristics of the visual system. For instance, insect compound eyes with larger facets have improved sensitivity due to increased light gathering ability, which in turn improves their ability to distinguish changes in contrast (Kirschfeld 1974). Conversely, smaller facets may reduce optical sensitivity but can improve spatial resolving power (Warrant and McIntyre 1993). Some insect eyes employ physiological strategies such as spatial summation where neurons in the lamina (the first optic ganglion in the insect brain) pool signals from the photoreceptors to improve sensitivity (Kirschfeld 1974; Warrant 1993; Nilsson and Ro 1994; Stöckl et al. 2016). Temporal summation is another physiological strategy whereby the integration time of the photoreceptors is extended to improve photon capture and enhance contrast discrimination, albeit at the cost of reduced temporal resolution (Frederiksen et al. 2008; Warrant 1999). These diverse and often conflicting strategies means that the visual system of an animal represents a trade-off between visual capabilities, such as sensitivity and spatial resolution, that are dictated by eye anatomy and physiology, and constrained by factors such as body size, lifestyle and visual ecology (Warrant and McIntyre 1993; Warrant 1999; Narendra et al., 2011; Somanathan et al. 2009b; Palavalli-Nettimi et al. 2019).

Bees are visually guided animals. Individual bees that exit their hive for the first time or from a newly discovered food source, perform exquisitely choregraphed learning flights, where they fly out, turn around and face the hive while retreating in a series of consecutive arcs that are roughly centred around the point of interest (e.g. Lehrer 1991; Zeil et al. 1996; Capaldi et al. 2000; Robert et al. 2016, 2018; Reyes et al. 2019; Collett and Hempel de Ibarra 2023). These flights are thought to allow bees to learn visual features around the goal which they rely on in subsequent trips (Cartwright and Collett 1987; Zeil and Wittmann 1993; Collett et al. 2013). Foraging bees must also cope with varying degrees of environmental ‘clutter’ and must employ strategies to avoid collisions with stationary and moving obstacles. Bees process the motion of images on their retina (i.e. optic flow) and translate this image motion into object range, thus relying on active vision strategies to perceive their three-dimensional world (Srinivasan et al. 1991; Hrncir et al. 2003; Eckles et al. 2012; Srinivasan 2021; Egelhaaf 2023). Bees use this information along with visuo-motor strategies, to control their flight speeds, stabilise gaze, detect obstacles and perform rapid flight manoeuvres to prevent collisions (e.g. Srinivasan and Zhang 2004; Boeddeker and Hemmi 2010; Baird and Dacke 2012; Burnett et al. 2020; Baird et al. 2021; Sathyakumar 2021; Ravi et al. 2022; Goyal et al. 2022). They use similar strategies to negotiate gaps and move safely between obstacles (e.g. Srinivasan et al. 1991; Baird and Dacke 2016; Ravi et al. 2019, 2020). In addition to being able to discriminate between colours (Giurfa et al. 1996; Hori et al. 2006; Spaethe et al. 2014; Hempel de Ibarra et al. 2014; Koethe et al. 2016), bees also recognize patterns and shapes that enables them to approach specific feeding sites or flowers (Dafni et al. 1997; Dyer et al. 2008, 2016; Hempel de Ibarra and Vorobyev 2009; Lunau et al. 2009; Sánchez and Vandame 2012). Back at their hive, bees use visual cues and visuo-motor strategies to defend their hive from intruders (Wittmann et al. 1990; Kelber and Zeil 1990; Koeniger et al. 2017). To carry out such behaviours, bees require a compound eye that has relatively high spatial resolution and good ability to detect contrast differences.

Several aspects of an insect’s visual physiology are correlated with size. Bees exhibit dramatic variation in body size varying from 1.8–38.0 mm in body length (Everaars 1979; Kelber and Somanathan 2019). Smaller bees tend to have smaller compound eyes—and thus fewer ommatidial lenses—which results in reduced spatial resolution and optical sensitivity (Kelber and Somanathan 2019, Ribi et al. 1989a; Spaethe and Chittka 2003; Somanathan et al. 2009a, b). In bumblebees, smaller workers have lower spatial resolution (interommatidial angles: 1.2° vertical and 2.9° horizontal), compared to larger workers (0.9° vertical and 2.1° horizontal; Spaethe and Chittka 2003). In other Hymenopterans, such as ants, species with fewer and smaller ommatidial lenses have reduced contrast sensitivity; however, their spatial resolving power is comparable to larger species (Palavalli-Nettimi et al., 2019).

Most research on the visual system of bees has focused on honeybees (*Apis*), bumblebees (*Bombus*), carpenter bees (*Xylocopa*) and sweat bees (*Megalopta*) (Ribi et al. 1989; Spaethe and Chittka 2003; Greiner et al. 2004; Somanathan et al. 2009b, a; Streinzer and Spaethe 2014). Stingless bees are one of the largest groups of bees (tribe: Melopinini), represented by over 550 species across the world (Grüter 2020). Spatial resolution has only been estimated in two stingless bee species: the Australian *Tetragonula carbonaria* and the South/South-East Asian *Tetragonula iridipennis*. Behavioural estimates of spatial resolution between the two species are comparable, at 0.053 cycles deg^-1^ (cpd) in *T. carbonaria* and 0.054 cpd in *T. iridipennis*. Anatomical estimates of spatial resolution in *T. carbonaria* are 0.32 cpd (Dyer et al. 2016), while theoretical estimates in *T. iridipennis*, are significantly lower at 0.17 cpd. Given this unexpected difference in *T. carbonaria* (behaviour vs anatomy estimates) and the variation between species (anatomical vs theoretical estimates), in this study we investigated spatial resolving power and contrast sensitivity of the compound eyes of *T. carbonaria* and *Austroplebeia australis* using pattern electroretinograms and compare this to the anatomical characteristics of the eye such as facet numbers and diameters.

Stingless bees have been an important part of indigenous Australian culture for centuries, far outdating modern meliponiculture which has become popular only in the last few decades (Heard and Dollin 2000; Halcroft 2012; Vit et al. 2013). Stingless bees such as *T. carbonaria* and *A. australis* are used as an important crop pollinator (Heard 1994, 1999; Heard and Dollin 1998; Vit et al. 2013). These two species are derived from phylogenetically different lineages (Rasmussen and Cameron 2009), that occupy overlapping ranges and exhibit distinct differences in their behaviour. *T. carbonaria* is a generalist and opportunistic pollen forager that invests more in resource collection, whereas *A. australis* tend to focus more on resource quality (Leonhardt et al. 2014). Additionally, *A. australis* foragers spend proportionately less amount of time hovering in front of flowers than *T. carbonaria*, hence are described as more ‘efficient foragers’ in terms of energy consumption (Halcroft 2012). *A. australis* are also active in slightly dim light conditions compared to *T. carbonaria* (Heard and Hendrikz 1993; Halcroft 2012). These two species therefore provide a unique opportunity to compare visual properties in a culturally and economically important species that forage in similar ecologies but differ in their foraging behaviour.

## Methods

### Study species

We investigated the compound eyes of two species of Australian native stingless bees: *Tetragonula carbonaria* Smith and *Austroplebeia australis* Friese (Fig. 1a). Experiments were carried out between January 2022 – February 2023, using worker bees collected from one hived colony of each species maintained at Macquarie University campus, North Ryde, NSW, Australia (33°46’10.24”S, 151°06’39.55” E).

**Figure 1:**
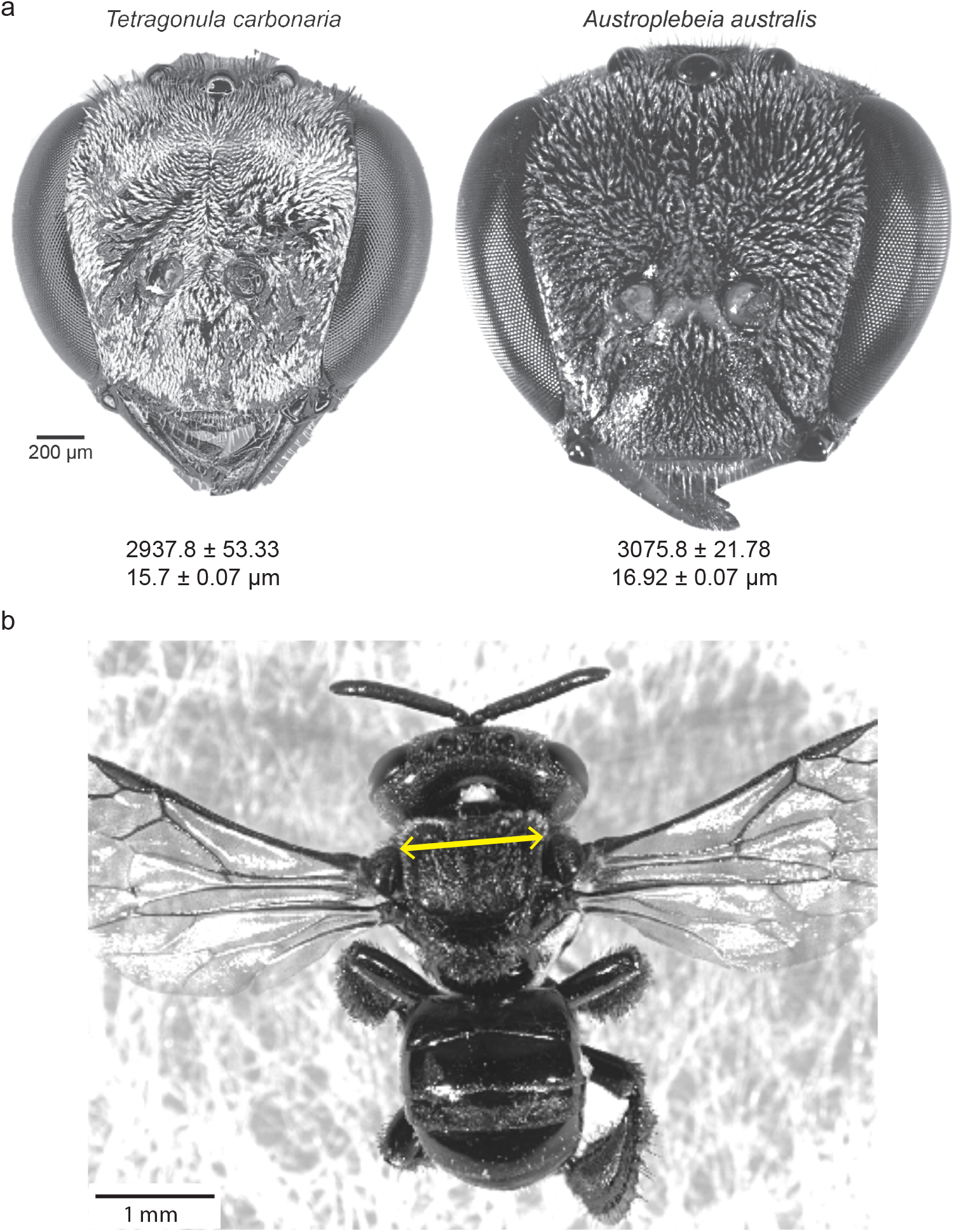
External morphology of stingless bee. **(a)** Gray scale dorsal view of the head of *Tetragonula carbonaria* and *Austroplebeia australis*. Average number of facets per eye (mean±s.e.m) and mean facet diameters of the medio-frontal regions of the compound eyes are shown. Image of *A. australis* courtesy of Giovanni Ramón-Cabrera. **(b)** Dorsal view of *T. carbonaria* showing intertegular width measurement, highlighted in yellow.

### External morphology of stingless bee eyes

We used the intertegular width (ITW), the distance between the wing bases, as an estimate of the body size of *A. australis* (n=5) and *T. carbonaria* (n=7; Fig. 1b). To determine the number of facets and facet diameters in the compound eyes of *A. australis* and *T. carbonaria*, we used an established technique described in detail elsewhere (Ribi et al. 1989; Ramirez-Esquivel et al. 2017). Briefly, we painted the surface of one eye with clear nail polish to create a cast of the facets. The cast was left to dry on the eye, removed and then flattened on a glass slide by making fine incisions. Eye casts were created for five individuals for both species and photographed under a light microscope (Leica DM5000B, Leica Microsystems GmbH, Wetzlar, Germany). For each individual, we counted the number of facets and measured the diameter of 260 randomly selected facets in the medio-frontal region of the compound eyes using ImageJ (version 1.52k) (Schneider et al. 2012). We determined the mean facet diameter of the mediofrontal region (the region with the largest facets) for each individual and from this we calculated the mean facet diameter for each species.

### Animal preparation for pattern electrophysiology

To investigate spatial vision in *A. australis* and *T. carbonaria*, we performed pattern electroretinograms (pERG). Bees were first anaesthetized by cooling them in an ice box for 5-7 mins before mounting them. To secure the bees, they were carefully manoeuvred into a 3 mm diameter pipette tip until the head and a small portion of the thorax was exposed. The pipette mount along with the bee was fixed horizontally onto a plastic stage with the bee’s dorsal side facing upwards using removable adhesive (Blu-Tack, Bostik Indonesia Inc., Banten, Indonesia). The head of the bee was oriented approximately 45 degrees from the mount to ensure that the compound eyes faced the stimulus, and the mandibles were fixed using beeswax to avoid movements. We removed the antennae in *T. carbonaria* but kept them in *A. australis* as this was the only way to ensure animals remained alive during the experiments. The animals were mounted within a Faraday cage wherein electrophysiological recordings were carried out. An active electrode of 0.25 mm diameter platinum wire with a sharp tip immersed in conductive gel (Livingstone International Pty Ltd., New South Wales, Australia) was positioned above the medio-frontal region of the compound eyes. A silver/silver chloride wire was inserted into the thorax of the bees, which served as an indifferent electrode, that was 0.1 mm diameter in *T. carbonaria* and 0.127 mm diameter in *A. australis*.

To reduce any effects of circadian rhythms on eye physiology, the experiments were conducted during the activity time of the species i.e., between 2 and 8 h post-sunrise.

### Pattern electroretinography (pERG)

The pERG technique was used to measure the spatial resolving power and contrast sensitivity of the compound eyes of *A. australis* and *T. carbonaria*. Detailed methods are described earlier (Ogawa et al. 2019; Palavalli-Nettimi et al. 2019; Ryan et al. 2020, Ogawa et al., 2023). Briefly, the experiments were carried out in a dark room within a Faraday cage. The bees were dark adapted for 10 mins prior to the first recording. We found this to be the maximum time that the bees could be dark-adapted for and remain alive for the duration of the experiment. The bees were then adapted to a uniform grey stimulus that had the same mean irradiance as the grating stimuli for 5 min.

The pERGs were amplified with a gain of x1000 and bandpass filtered between 1 Hz and 1 kHz with a differential amplifier (DAM50, World Precision Instruments Inc., FL, USA). Voltages were digitised at 20000Hz using a National Instruments data acquisition device (USB-6353, National Instruments, Austin, TX, USA) controlled via custom software written in Microsoft Visual Studio (2013, Microsoft Corporation, Redmond, WA, US) which was also used to control the stimulus presentation.

Visual stimuli were projected by a digital light processing (DLP) projector (W1210ST, BenQ corporation, Taipei, Taiwan) onto a white screen (51 cm width × 81 cm height) at a distance of 30 cm from the animal. We used vertical contrast-reversing sinusoidal gratings of 11 spatial frequencies (0.6, 0.5, 0.45, 0.4, 0.35, 0.3, 0.25, 0.2, 0.15, 0.1 and 0.05 cpd) and up to eight Michelson’s contrasts contrasts (95%, 85%, 75%, 50%, 25%, 12.5%, 6% and 3%) for each spatial frequency in descending order. Every second spatial frequency was presented in descending order, followed by the alternative spatial frequencies in ascending order to assess if signal strength degraded over time. As a control, the non-visual electrical signal (background noise) was recorded at two spatial frequencies (0.05 and 0.1 cpd) at 95% contrast with a blackboard to shield the animal from the visual stimuli before and after running the experimental series. The maximum signal out of four control runs was used as the noise threshold.

The sinusoidal gratings were reversed at a temporal frequency of 2 Hz. For each combination of the stimuli, 10 repetitions of the response for 5 s each were averaged in the time domain. The averaged responses were then analysed using a Fast Fourier Transform, FFT. An F-test was used to assess whether the response signal at the second harmonic (4 Hz) of the FFT response spectrum differed significantly from 10 neighbouring frequencies, five on either side, for each presented stimulus. Spatial resolving power and contrast threshold were obtained by interpolating from the last point above the noise threshold whose FFT amplitude at 4 Hz was also significantly greater than the 10 surrounding frequencies, and the first point below the noise threshold. Contrast sensitivity is defined as the inverse of contrast threshold.

## Statistical Analyses

We performed a one-way ANOVA in R (R Core Team 2018) to identify if the facet diameters and facet numbers varied between *T. carbonaria* and *A. australis*. A linear model in R (R Core Team 2018) was used to test if the maximum contrast sensitivities and spatial resolving powers were different between the two species. Subsequently, we used a linear mixed-effects model to investigate whether the spatial frequency of the pERG stimulus and the species affected contrast sensitivity functions. Spatial frequency of the stimulus and the species were the fixed effects while the animal identity was a random effect. We carried out a pairwise comparison to identify the spatial frequencies at which the contrast sensitivities of the two species were different using the *emmeans* package in R (R Core Team 2018), with a Bonferroni correction applied.

All linear mixed effect models were carried out in the lme4 package (Bates et al., 2015) of R (R Core Team 2018) using *lmer* with the restricted maximum likelihood (REML) estimation method. The Akaike Information Criterion function in R (R Core Team 2018) was used to find the best fit for all regression models and a value of 10 was used as the cut-off. The significance of the fixed effect terms was examined using the t-test with Satterthwaite approximation for degree of freedom (lmerTest package). In all instances mentioned above, the model assumptions such as linearity were tested by plotting the residuals against the fitted values of the model.

## Results

### External morphology of A. australis and T. carbonaria eyes

The mean ITW of *A. australis* was measured as 1.27 ± 0.02 mm (mean ± s.e.m), while that of *T. carbonaria* was 1.15 ± 0.02 mm (Fig. 1b; Table 1). Facet number varied significantly between the two species, with approx. 5% more facets in *A. australis* (3075.8 ± 21.78; mean ± s.e.m) compared to *T. carbonaria* (2937.8 ± 53.33; one-way ANOVA: F_(1,8)_=5.74, p=0.04). Facet diameters also varied between the two species with larger facets in *A. australis* (range: 13.2–22.51 μm) compared to *T. carbonaria* (11.04–19.69 μm; Fig. 2; Table 1; *F*_*(1,8)*_*=6*.*15, p=0*.*04, One-way ANOVA*).

**Table 1.** External morphology and visual properties of Australian stingless bees and European honeybee. Data for *A. mellifera* from Somanathan et al. 2009b* and Ryan et al. 2020^†^.

| Measurement<br>(mean ± s.e.m) | Species |  |  |
| --- | --- | --- | --- |
|  | *A. australis* | *T. carbonaria* | *A. mellifera* (n=5) |
| Intertegular width (mm) | 1.27 ± 0.02<br>(n=5) | 1.15 ± 0.02<br>(n=7) | 3.2* |
| Mean number of facets | 3075.8 ± 21.78<br>(n=5) | 2937.8 ± 53.33<br>(n=5) | 5311 ± 186<sup>†</sup><br>(n=5) |
| Mean facet diameter (µm) | 13.2 – 22.51<br>(minimum-maximum) | 11.04 – 19.69<br>(minimum-maximum) | 18.8 ± 0.6<sup>†</sup><br>(30 facets/individual) |
| Spatial resolution (cpd) | 0.53 ± 0.03<br>(n=5) | 0.53±0.02<br>(n=5) | 0.54 ± 0.02<sup>†</sup><br>(n=5) |
| Maximum contrast<br>sensitivity (at 0.05 cpd) | 13.07 ± 1.17 (7.65%)<br>(n=5) | 5.99 ± 1.09 (16.68%)<br>(n=5) | 16.92 ± 3.0<sup>†</sup> (6%)<br>(n=5) |

**Figure 2:**
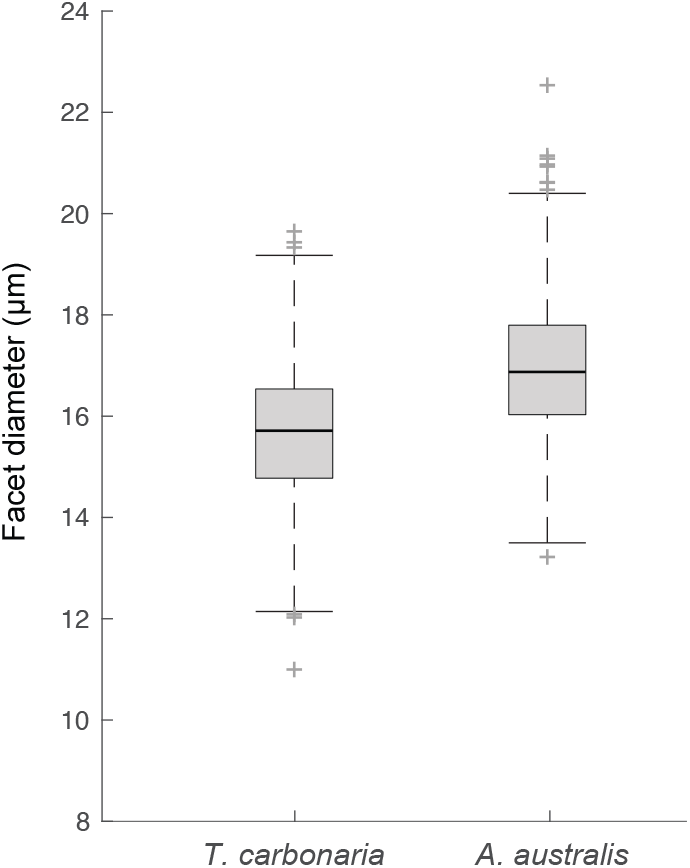
Facet diameters of stingless bees. Boxplots showing all the measured facet diameters of *T. carbonaria* and *A. australis*. Plus (+) symbol indicate outliers.

### Spatial resolving power and contrast sensitivity of A. australis and T. carbonaria

The spatial resolving power of the two species did not differ (*A. australis*: 0.53 ± 0.03 cpd; mean ± s.e.m; *T. carbonaria*: 0.53 ± 0.02; Fig. 3b; Table 1; *Linear model*: *F*_*(1,8)*_*=0*.*002, p=0*.*96*). As the spatial frequencies of the stimulus increased, the contrast sensitivity decreased in both species (Fig. 3a; Table 2). The maximum contrast sensitivity attainted at the lowest spatial frequency (0.05 cpd) was significantly higher in *A. australis* at 13.07 ± 1.17 (7.65%; mean ± s.e.m) than in *T. carbonaria* at 5.99 ± 1.09 (16.68%; Fig. 3a; Table 1; *Linear model*: *F*_*(1,8)*_*=11*.*42*; *p=0*.*009*). In *A. australis*, the contrast sensitivity was significantly higher at all spatial frequencies except at 0.26 and 0.31 cpd (Fig. 3a; Table 2; *T. carbonaria*: 0.23 cpd, *A. australis*: 0.23 cpd; *p=0*.*006*).

**Table 2.** Summary of linear mixed-effects model to test the relationship between the contrast sensitivity function, spatial frequency of the grating stimulus and the species tested. Model: contrast sensitivity ∼ spatial frequency * species +(1 | individual identity). The t-tests for fixed effects use Satterthwaite approximations to degrees of freedom.

| Parameter | *Estimate* | *Standard Error* | *Degrees of freedom* | *t-value* | *p-value* |
| --- | --- | --- | --- | --- | --- |
| Intercept | -0.09 | 0.04 | 53.59 | -2.47 | 0.02 |
| Spatial frequency | -0.66 | 0.05 | 83.09 | -14.64 | $< 2e^{-16}$ |
| *A. australis* | -0.06 | 0.05 | 47.11 | -1.16 | 0.25 |
| Spatial frequency: *A. australis* | -0.26 | 0.06 | 83.26 | -4.22 | $6.31e^{-5}$ |

**Figure 3.**
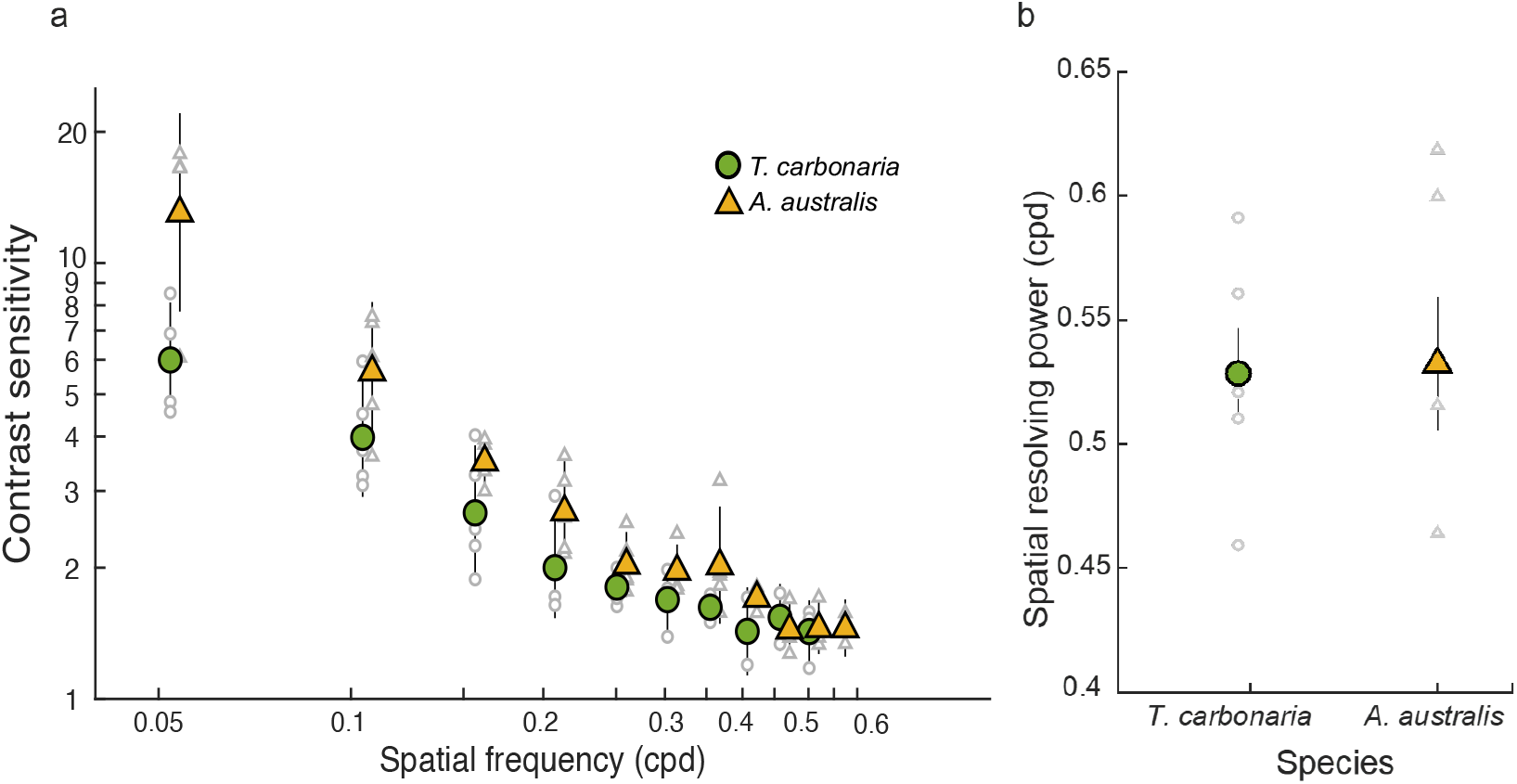
Contrast sensitivity (a) and spatial resolving power (b) of the compound eyes of the stingless bees *T. carbonaria* and *A. australis* (n=5 for both species). In (a) each coloured data point is the mean contrast sensitivity of all individuals of a particular species at the corresponding spatial frequency. The error bars show 95% confidence intervals. Individual data points are shown in grey. Data points for *A. australis* were shifted to the right of the recorded spatial frequency to improve visualisation. All contrast sensitivity values plotted on a log scale. In (b) each coloured data point is the mean spatial resolving power of all individuals of a particular species at 95% contrast. Error bars (black lines) show standard error (s.e.m.). Individual data points are shown in grey.

## Discussion

We investigated the spatial properties of the compound eyes in two Australian stingless bee species. The larger species, *A. australis*, had more facets and larger facets compared to the smaller *T. carbonaria*. The spatial resolving power of the two species did not differ significantly; however, the larger *A. australis*, had higher contrast sensitivity compared to the smaller *T. carbonaria*.

### Spatial resolving power of compound eyes of bees

Compared to the two species of stingless bee studies here, the honeybee *A. mellifera* has almost 1.5 times more facets. However, the spatial resolving power of both stingless bee species was 0.53 cpd (Fig. 3b, Table 1) and this is comparable to *A. mellifera* (0.54 cpd; Table 1; Ryan et al. 2020). The methods used to study stingless bees and honeybees were identical, which makes these results directly comparable. By comparison, in ants, where the same pERG technique was used, spatial resolving power decreased with the number of facets, although this relationship was not linear. The smallest ants that had only 8% of the facets compared to the large-eyed ants, still had 80% of the spatial resolution compared to the largest animals (Palavalli-Nettimi et al. 2019). A similar trend appears among the bees in this study. Bees, owing to their flying mode of locomotion have larger eyes and more facets per eye compared to worker ants that lead a pedestrian lifestyle. It is interesting that species with such differences in their external sensory array and lifestyles have similar spatial resolution. The variation in eye anatomy of bees is important, as it influences their contrast sensitivity which is maximised to suit their lifestyle and body size limitations.

Bees of the genus *Tetragonula* are the only stingless bees in which spatial resolution has been estimated using theoretical techniques (e.g., μCT, tracking pseudopupil changes) and behavioural methods (single target detection ability) (Kelber and Somanathan 2019). The theoretical estimate of spatial resolution of *T. iridipennis* is 0.17 cpd, and the behavioural estimate is 0.04 cpd (chromatic contrast only) and 0.054 cpd (chromatic + achromatic contrast). The theoretical estimate of spatial resolution of *T. carbonaria* is 0.3 cpd (e.g., pseudopupil changes), and the behavioural estimate is 0.053 cpd (both chromatic only and chromatic + achromatic contrast). Behavioural and theoretical estimates of spatial acuity typically do not match in *A. mellifera* (Ryan et al. 2020). The discrepancies in the estimates of spatial resolving power from pseudopupil illumination and the pERG are potentially due to the differences in the technique. Pseudopupil tracking method measures the axial directions of ommatidia whereas pERG relies on extracellular signals from the lamina where both temporal and spatial summation strategies may occur. The behavioural estimates of spatial acuity in *T. carbonaria* suggest that bees detect a 1 cm diameter flower only when they were 6 cm from the target. Although, in natural outdoor experiments, bees detect thin silk strands of spider webs at this distance (Sathyakumar 2021), which likely requires a higher behavioural spatial acuity. It would be thus useful to assess their spatial resolution using behaviours which rely on high spatial resolution such as object detection or avoidance.

### Contrast sensitivity of compound eyes of bees

Contrast sensitivity is the ability to discriminate patterns as their brightness contrast decreases (Land 1997). pERG measurements showed that at the lowest spatial frequency of 0.05 cpd the maximum contrast sensitivity was 13.07 (7.65% contrast) in *A. australis* and 5.99 (16.68%) in *T. carbonaria* (Fig. 3a, Table 1). In addition to having higher contrast sensitivity, *A. australis* bees also have more and larger facets (Figs. 1a, 2, Table 1). At present, we lack anatomical data on these stingless bee species. If the rhabdoms of *A. australis* are wider and the focal length is shorter compared to *T. carbonaria*, it would account for their increased optical sensitivity and improved contrast sensitivity. Indeed, day-active bees (*Tetragonula iridipennis*) and ants (*Temnothorax rugatulus*), with short focal lengths have higher than expected optical sensitivity that suggests they may also have higher contrast sensitivity (Ramirez-Esquivel et al. 2017a; Jezeera et al. 2021). So, why do *A. australis* require higher contrast sensitivity compared to *T. carbonaria*? While there are no studies that have specifically addressed this, there are a few possibilities based on behavioural observations. *A. australis* are active at low light intensities of up to 5.8 W m^-2^ (Halcroft 2012) compared to *T. carbonaria* whose activity stops at 15 W m^-2^ (Heard and Hendrikz 1993). Other Hymenoptera such as nocturnal ants that forage in low light also tend to have increased contrast sensitivity (Ogawa et al. 2019).

Interestingly, *A. australis* exhibited similar contrast sensitivity to *A. mellifera* despite having significantly fewer facets and being considerable smaller in body size (Ryan et al. 2020). This suggests that good contrast sensitivity is relatively important for *A. australis*, and perhaps other visual tasks such as colour discrimination or temporal resolution may be more important for *A. Melifera* and *T. carbonariai*. A higher contrast sensitivity could enable *A. australis* to discriminate between many different shades and colours of petals. There is some indication from analyses of the collected pollen that *A. australis* collect pollen from a relatively narrow flower colour spectrum compared to *T. carbonaria* (Leonhardt et al. 2014). Better contrast sensitivity perhaps combined with spectral sensitivity could allow these *A. australis* to detect spatial patterns more efficiently compared to *T. carbonaria*. This remains to be tested since we still know very little about the ecology and behaviour of native bees.

In conclusion, we found that the Australian native stingless bee *A. australis* have more and larger facets and higher contrast sensitivity in their compound eyes compared to *T. carbonaria*. These differences did not affect their spatial resolving power, which was relatively high and comparable to the much larger European honeybee, indicating the significance of spatial resolution in these pollinators. Now that we know the physiological capacities of the compound eyes of stingless bees, it sets the scene to investigate the behavioural limits of visually guided behaviours in these important pollinator species.

## Acknowledgements

We are grateful to the Australian native bee research community and members of the Australian Native Bee Association (ANBA) for their enthusiasm and for their time to discuss different aspects of this project. We thank Rosalyn Gloag and Dan Smailes for providing us with a hive of *A. australis*. We acknowledge the Wallumattagal clan of the Dharug nation as the traditional custodians of the Macquarie University land where we conducted our research. We collected specimens from the lands of the Dharug people.

## Diversity and Inclusion

We strongly support equity, diversity and inclusion in science. The authors come from different countries (India, Japan, Australia and the United Kingdom) and represent different career stages (PhD candidate, postdoctoral researcher, senior lecturer, professor). Three of the authors are from underrepresented ethnic minorities in science. Three of the authors self-identify as female, an underrepresented gender in science.

## Author contributions

Study design: BP, AN; Data collection and analyses: BP, LR; Built the equipment and wrote the software: NSH, LR, YO; Writing – original draft: BP; Writing – review and editing: all authors; Visualisation: BP, AN; Funding acquisition: AN.

